# A systems biology investigation to identify potential biomarkers in leprosy-affected host cells

**DOI:** 10.1101/2025.03.28.646010

**Authors:** Shomeek Chowdhury, Afzal Ansari, Pranshi Verma, Anuj Mavlankar, Anjali Tripathi, Maria Pena, Rahul Sharma, John Caskey, Richard Truman, Pushpendra Singh

**Author notes:** These authors contributed equally to this work (Shomeek Chowdhury, Afzal Ansari).

## Abstract

*M. leprae* adapts to the host cell environment and disrupts protein-protein interactions inside the host cell for its own survival and proliferation. This study is attempts to comprehend these host-pathogen interactions at the systems level. We utilised RNA-Seq data set of leprosy-resistant and leprosy-susceptible armadillos and investigated how five different human signaling pathways get hijacked by *M. leprae.* We also identified the corresponding human homolog proteins. By applying graph theory on these protein networks, we predicted 25 proteins which play important role in these pathways related to leprosy pathogenesis and/progression. The study is also supported by wet lab experiments identifying 69 proteins in armadillos upon leprosy progression after experimental infection which get differentially expressed. Our study found that 69.5% of these experimentally identified proteins are part of either *M. leprae*-affected signaling pathways or possibly contribute to leprosy pathogenesis based on their GO terms. This study identified three human proteins: LC3-II, PI3K and IDO1 which have been implicated in *M. leprae* pathogenesis in other studies also leprosy patients. Thus, upon further experimentally validation, such markers can be potentially useful to diagnose leprosy in patients at an early stage .

## 1. Introduction

Leprosy in last decade has shown a rapid decline in the global prevalence, but, still remains a significant health problem in endemic countries, such as India which records more case than the rest of the world [1]. Leprosy diagnosis has till date focused on the appearance of clinical symptoms for example, lesions and loss of sensation which can gradually lead to nerve damage if not detected and treated timely with multidrug therapy (MDT) [2,3]. Therefore, delayed detection leads to problems, and there should be proper strategies for early detection of the disease. *M. leprae* has the capability of creating a favorable environment for its own inside human host cells. Like other pathogens, this bacterium also disrupts the host signaling interactions [4–8]. But the strategies which it makes to do so are very unique and these need to be investigated further to gain better understanding leprosy pathogenesis and in order to determine the host proteins which are regulated by it. Identification of these host proteins playing major roles during leprosy can be designated as host biomarkers which can be later used for diagnostic purposes for early detection of the disease [9–12]. Experimental investigations for determining host biomarker proteins requires study of protein-protein interactions (PPI) which requires significant amount of resources and skills [13,14]. Such investigations need to study the relationships between different pathways and their interactions i.e. how they are connected. Systems biology and computational modeling offer great promise for answering these questions. Identification of such dysregulated pathways and their associated interactions will help us in predicting potential biomarkers which can be prioritized for experimental validation using wet-lab tools [15–17].

There is a need to understand the signaling routes followed by *M. leprae* which it uses to propagate itself and cause disease progression in humans. It is a very well-known fact that most of the pathogens once they enter the body, affect the crucial host signaling pathways to disrupt the protein-protein interactions by inhibiting a protein activity or causing excessive activation of another protein [8]. From the list of genes identified in this study, progression of leprosy is affecting five major host signaling pathways: mTOR, TLR, T-cell receptor (TCR), MAPK and NOTCH [4,18–21]. It is true that *M. leprae* related immunological processes might be involved in additional pathways. It has actually been seen in JAKSTAT signaling pathway and NF-kB pathways [22,23]. But these and many other small signaling networks are already part of the five global pathways studied in this work using computational biology approaches. These subset pathways have been included in the five signaling networks, for example, JAK-STAT and NF-kB pathways are part of T-cell, TLR and NOTCH signaling networks. Also, the literature articles (cited in this manuscript later) showed these five networks to be affected which is consistent with our results. Hence, we have chosen these five pathways. The influence of it over these big cascades can help us to realize the extent to which this organism is regulating the host cellular environment. In mTOR, *M. leprae* enters the host cell in a phagosome. ESX-1 inside the host cell is released by the bacteria. This protein frees the bacteria from the phagosome and then, using its coat protein, LEPLAM, bacteria interact with PI3K & CALMODULIN to inhibit the signaling through these proteins as PI3K along with other proteins causing the down-regulation of the *M. leprae* activities in the cell [24,25]. LC3-II alters these dynamics hugely whose main function is to form the autophagosome around the bacteria to kill it [26,27]. *M. leprae* proteins LPRG, BFR, and PGL-1 make many interactions with the TLR pathway proteins [28–30]. TLR1/2 and TLR4 are the signaling cascades which are used by the bacteria for evasion of host immunity [19,31]. Additionally, TLR6 and TLR9 cascades are also used by the bacteria to spread the disease [32,33]. Nitric oxide (NO) production from INOS and IL12 production from NF-kB is inhibited during leprosy progression through TLR pathway as these are known to secrete cytokines to stimulate the required immunological response which is not good for the bacteria [32,34,35]. T-cell signaling pathway is used by *M. leprae* for infection through the MHC-CLASS-II proteins. LPRG and BFR are recognized by MHC CLASS II which stimulates the T-cell pathway to produce inflammatory cytokines IL12, IFN GAMMA and IL2 etc [29,34,36,37]. Four *M. leprae* proteins are involved in the MAPK pathway which are ML-LBP21, PGL1, LEPLAM, MURAMYL DIPEPTIDE and PIM2 [30,38–41]. ERK1/2 acts as the central protein in the leprosy affected MAPK pathway [42]. In NOTCH pathway, BFR, LPRG, PIM2 and PGL1 are the 4 proteins produced by *M. leprae* [28,29,41,43]. NOTCH1 and NOTCH2 signaling are hugely involved in leprosy with CSL as the main transcription factor (TF) [44]. The entire NOTCH signaling occurs during immunopathogenesis of leprosy where the output proteins HES1 and SOCS3 are produced [45].

It is important to identify the proteins which are playing crucial roles during leprosy in the aforementioned five signaling pathways which can be later designated as biomarkers for diagnostic or prognostic purposes. We have used two terms called “Over Activity” and “Under Activity” for the proteins getting up-regulated and down-regulated respectively according to our computational results. Pathway reconstruction through proper literature survey is important for obtaining a broader picture of cellular processes and therefore we have considered this methodology for this work [46,47]. We, for the first time, have constructed leprosy based signaling pathways where we have tried to incorporate maximum number of interactions from literature between the bacteria and the host proteins [48]. Using databases and literature surveys, we are till date claiming the most comprehensive global pathway maps for the five *M. leprae* infected signaling networks.

Looking at these big systems in their entirety using experimental tools is not feasible, so computational modeling approaches for investigating them can give us understanding of the intricate and complex mechanisms of the system in a short period of time. Hence, graph theoretical analysis was used in this study to identify the biomarkers or the host proteins in the network which are getting hugely regulated. Graph theory for analyzing biological networks is a regular phenomenon in the field of systems biology [49,50]. This paper explains the methodologies of graph theory used to investigate biological networks. Graph theory has already been used to understand the disease mechanisms through PPI networks. Safari-Alighiarloo et al suggested that these methods help in understanding the pathogenic mechanism which causes the progression of disease [51]. Spirin et al has mentioned that graph theory can give us the important proteins in a network like splicing or transcription factors [52]. Not only molecular networks, graph theory nowadays are also being used to identify the target proteins of a drug [47,53]. These evidence show that despite being a qualitative approach, graph theory or network analysis has been regularly applied on robust biological systems to comprehend the biological dynamics in a real fashion. Knowledge based or literature mining also has been simultaneously used for graph theoretical approaches which this paper has depicted [54].

Experimental works in the field of leprosy till date has been done extensively. Geluk et al have identified biomarkers in blood samples of leprosy patients [12]. Also, *Mycobacterium indicus pranii* (MIP) shares antigen with *M. leprae* and the proteins regulated by this organism in the host cells are very much similar to what *M. leprae* regulates which is evident from the biomarkers we found in our analysis [55]. Computational analysis works related to this disease are very few. However, people have tried to investigate the biological dynamics using computational modeling for other pathogenic diseases like leishmaniasis and tuberculosis [56,57]. But no one till date has tried to investigate the signalling pathways affected in leprosy progression It is noteworthy that there are some similarities in leprosy and tuberculosis pathogenesis (i.e. granuloma formation etc) [25]. In order to study the incidence and dissemination of leprosy and for proposing diagnostic strategies of this disease, we have to understand this biological process using computational modeling. Present study has investigated these five host signaling pathways mentioned in detail using experimental evidence in literature based upon PPI in leprosy pathogenesis related publications [18,20,21,31,37]. We then performed the simple qualitative approach graph theoretical analysis to identify biomarkers or the proteins which are regulated during the disease [17,49]. The biomarkers are again validated from literature. In addition to that, graph theory of it gave us important host biomarkers which are crucially regulating the disease and can be later on used as diagnostic biomarkers [11,12,53].

## 2. Materials and Methods

### 2.1 Experimental setup

Armadillos were captured from the wild and acclimated to the laboratory environment following established protocols [58,59]. To rule out any pre-existing natural infection with *M. leprae*, the animals were screened for *M. leprae*-specific antibodies over a period of six months before experimental infection. The study received ethical approval and was conducted in accordance with the U.S. Public Health Service policy on the care and use of laboratory animals (NHDP IACUC assurance number A3032-01).

10 armadillos were challenged with high-dose experimental infection of live *M. leprae* intravenously and their Peripheral blood mononuclear cells (PBMCs) were collected and cryo-preserved using liquid nitrogen at the 4th and 18th month time-points post-infection. Animals were monitored for the *M. leprae* specific antibody titers using ELISA and nerve conduction velocity. The PBMCs were thawed and plated for further stimulation with various *M. leprae* antigens. RNA was extracted, followed by library preparation for RNA-Sequencing. An analysis pipeline assessed differential expressed genes (DEGs) between susceptible and resistant animals.

### 2.2 Pathway Reconstruction from databases and literature

First in order to get an idea, the pathway databases were searched to get the big picture of the five signaling networks like Reactome, Cell Signaling Technology, Pathway Central, Sigma Aldrich, KEGG and Protein Lounge [60–63]. Databases from time to time have been subjected to discrepancies; hence, there is a need to validate the interactions from literature. So, for all the interactions we got from the database, we again did the literature search for that interaction. For most of the interactions, we tried to find those papers where the protein interaction is being showed during or is related to leprosy in some or the other way. We created the interaction list with PUBMED reference for each of the protein-protein interaction for each of the five signaling pathways [64].

### 2.3 Experimental Proteins Inclusion into the pathways

Once, the experimental investigation was done in the form of RNA sequencing of leprosy affected armadillos, the transcriptomic profiles were analyzed in both susceptible and resistant armadillos, which identified 81 differentially expressed genes (DEGs) in this organism during leprosy progression. Out of these 81, we got 69 proteins whose human homologs have been found. As these proteins are involved in leprosy, we hypothesized that they should also be engaged in the leprosy-affected networks. From interaction database BIOGRID and from literature articles where protein interactions are given on the basis of yeast two hybrid experiments [65], we have tried to find the interaction partners of these 69 proteins and the cascade through which they are involved in leprosy [49]. Finally, we could map 48 of these proteins in these five signaling pathways. Using Gene Ontology (GO) biological process terms, we have also tried to find the enrichment of these 48 proteins in the two most common and broad biological processes related to leprosy, immunological and neurological mechanisms [50,51].

### 2.4 Pathway drawing in Cell Designer

The five leprosy affected human signaling pathways, NOTCH, T-cell, MAPK, mTOR and TLR were drawn in the pathway visualization software called Cell Designer [66]. This software follows programming language Systems Biology Markup Language (SBML) which is the most appropriate computing platform for these systems biology studies [67]. This software provides legends and connectors to denote the protein, enzymes, and activating/inhibiting interactions in a form which makes the biological pathways clearly understandable.

### 2.5 Gene Ontology (GO) Biological Process enrichment for human homologues protein

The 69 proteins were subjected to Gene Ontology Biological Process enrichment using the Uniprot interface [68,69]. The biological process for each of the 69 proteins were checked and were divided into two types: immunological functions and neurological functions as these are the functions in which leprosy related activities are performed [2,27].

### 2.6 Graph theoretical modeling of the signaling pathways

Gephi software was used to perform the graph theoretical modeling of the five signaling networks [70]. The six topological measures we used to perform the network modeling of the signaling networks are defined below [9,47,49].

**In-degree:** It refers to the total number of edges or connections (activations or inhibitions) acted by different nodes of a network on a particular node.

**Out-degree:** It refers to the total number of edges or connections (activations or inhibitions) acted on a particular node by other nodes in the network.

**Total-degree:** It refers to the total number of in-degree and out-degree of a particular node.

**Eigen vector centrality:** It refers that a node in a network will be more central if it is connected to many central nodes in the network.

**Betweenness centrality:** It is the ratio of the number of shortest paths (**minimum number of intermediate links or connections that has to traverse from on node to other node in the network**) that pass through the node to the total number of shortest paths of all the nodes to all the other nodes.

**Closeness centrality:** It is defined as the inverse of sum of the total length of the distances or shortest paths of one node to other nodes in the network.

## 3. Results

### 3.1 Experimental results

RNA-Seq results of 10 armadillos (n=6 resistant and 4 susceptible) were compared after normalization for differential gene expression and hierarchical clustering using standard bioinformatics and statistics tools (Figure 1). The most differentially expressed genes between the two groups of animals showed distinct clustering of resistant and susceptible animals except in one case where a susceptible animal was found to cluster with the group of resistant animals. This analysis gave us 81 genes in armadillos whose human homologs have been found which were found to be significantly up-regulated/down-regulated in the animals susceptible to leprosy. Out of the 81 genes, the ENSEMBL Gene ID’s were present for just 69 proteins as for the other 12, matching ID’s were not found [71]. With these 69 host proteins obtained by RNA Sequencing, we started the analysis of computational validation of these experimental results (Table S3).

**Figure 1.**
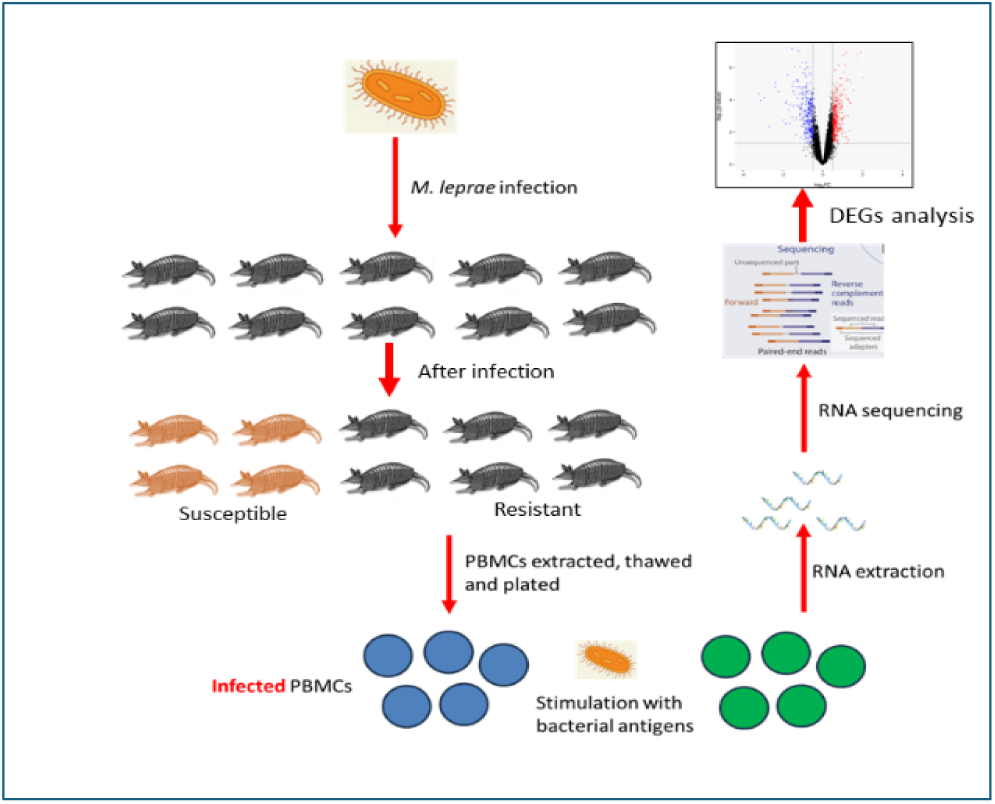
**Wet lab protocol for obtaining differentially expressed genes following *M. leprae* infection.**

**Figure 2.**
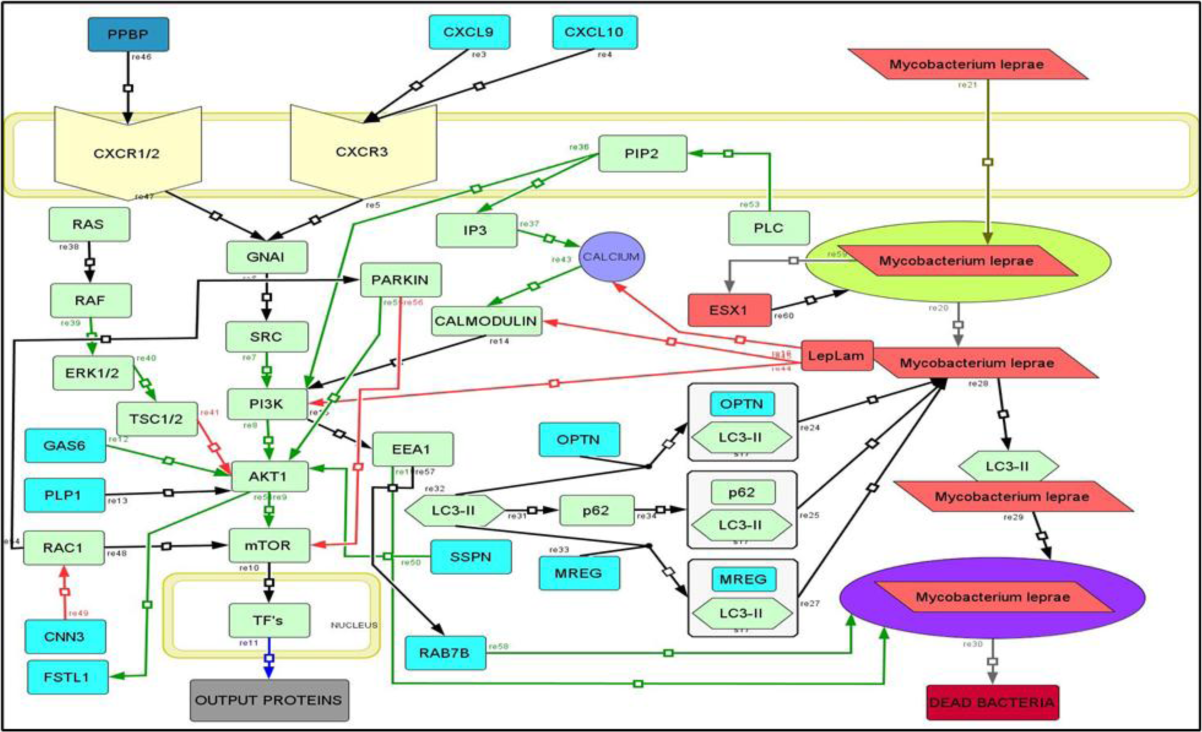
Leprosy-affected mTOR signaling pathway. Protein LC3-II form complexes with various host proteins to facilitate killing of M. leprae via autophagosome formation. M. leprae protein LEPLAM inhibits calcium and Pi3K-AKT signaling directly to take control over the host cell. Downregulated DEG proteins are shown in blue and other host proteins are shown in green. *M. leprae* proteins are shown in red. Physical interaction, Activation, Inhibition, Protein production, Protein Release and Membrane Translocation are shown using black, green, red, blue, grey and olive-green arrows.

**Figure 3.**
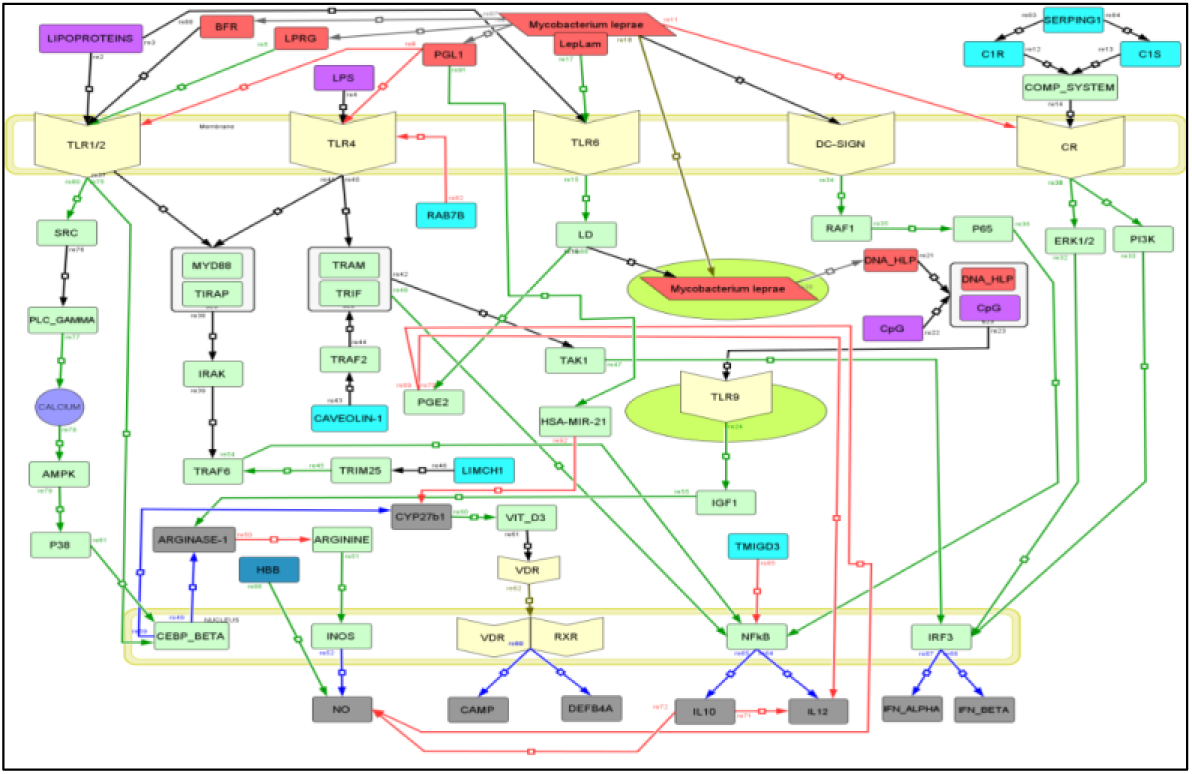
*M. leprae* adapting to host cell environment via TLR signaling pathway. *M. leprae* protein DNA-HLP by forming complex with human CpG islands bind TLR9 receptor which leads to activation of ARGINASE-1 protein. This enzyme cleaves arginine which *M. leprae* wants as arginine produces nitric oxide (NO) which inhibits *M. leprae-*host protein interactions. Downregulated DEG proteins are shown in blue and other host proteins are shown in green. *M. leprae* proteins are shown in red. Physical interaction, Activation, Inhibition, Protein production, Protein Release and Membrane Translocation are shown using black, green, red, blue, grey and olive-green arrows.

**Figure 4.**
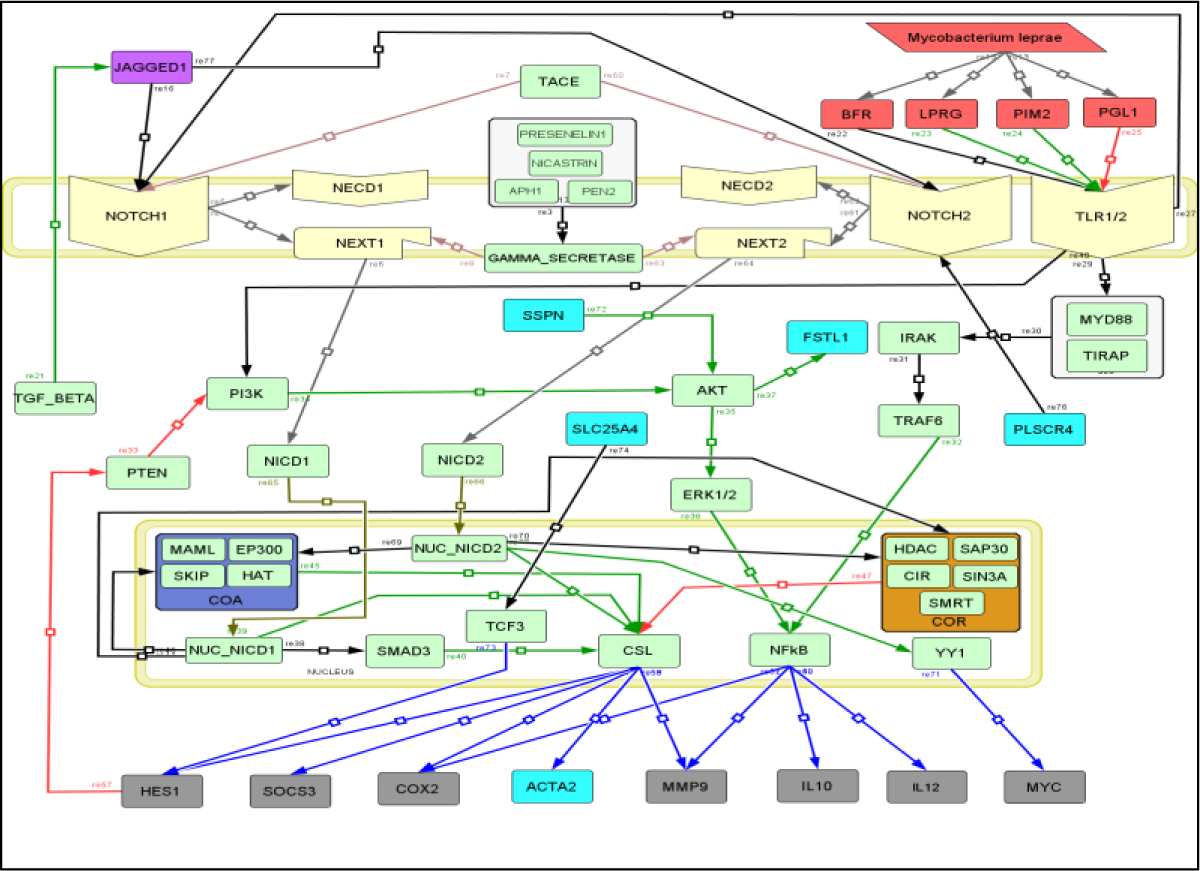
*M. leprae* can hijack NOTCH signaling pathway. *M. leprae* proteins interact with TLR receptors which activate signaling routes via 1) PI3K-AKT and 2) MYD88-IRAK. Both these signaling cascades lead to activation of transcription factor NF-kB which produces ACTA2 (one of the differentially expressed proteins during leprosy).

**Figure 5.**
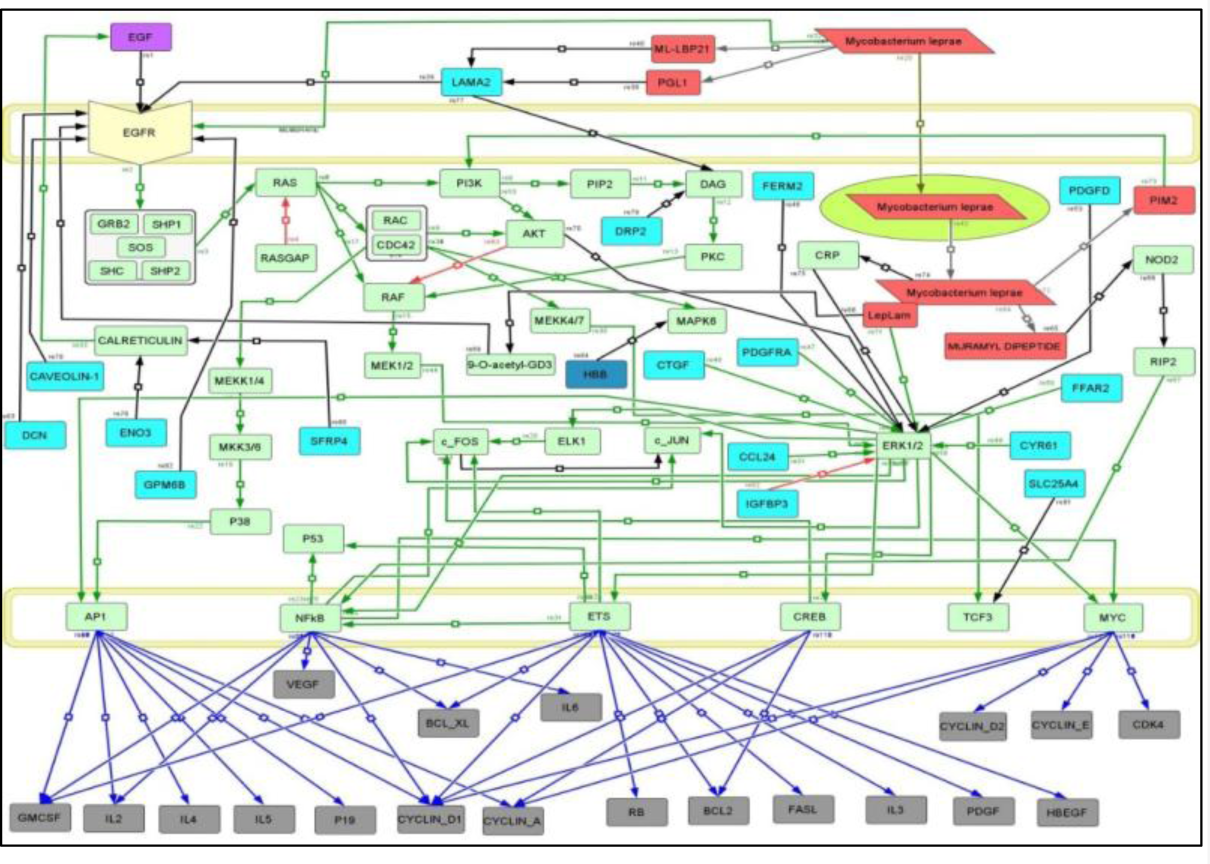
MAPK signaling pathway during leprosy. Multiple host proteins active during leprosy (differentially expressed) and *M. leprae* proteins make protein-protein interactions with receptor EGFR indicating crucial functioning of MAPK pathway in leprosy progression.

**Figure 6.**
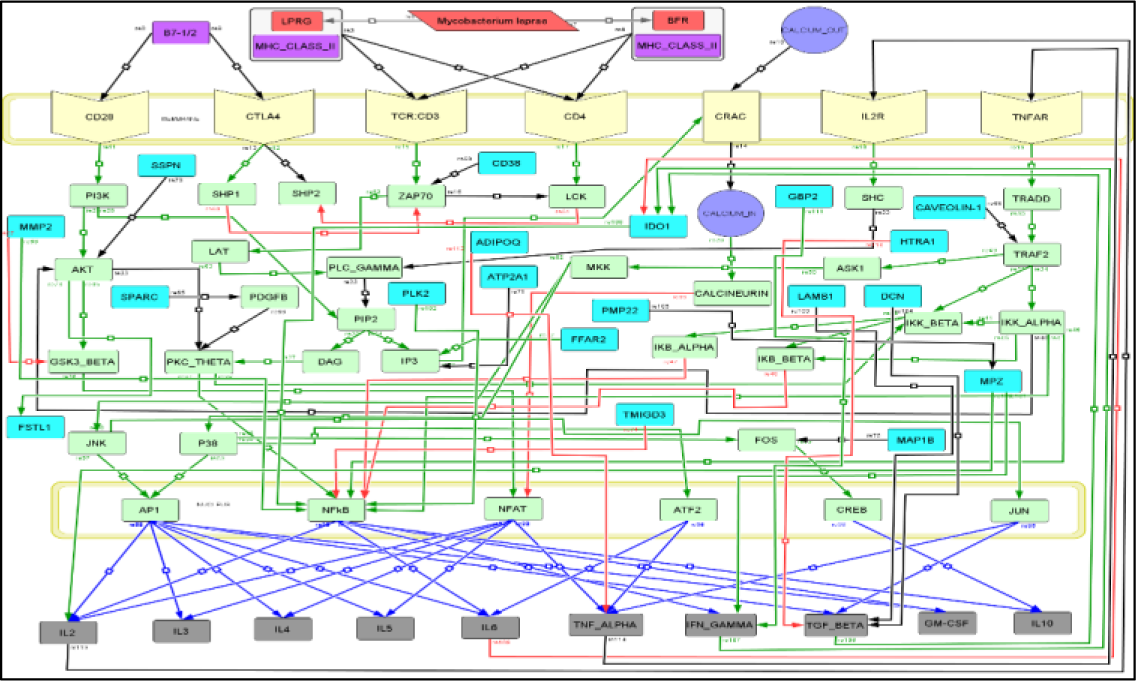
*M. leprae* disrupts T-cell signaling interactions to cause leprosy. Once the bacterial antigen is presented by MHC complex to T-cell, specific protein-protein interactions of T-cell affected leading to high production of proteins such as IFN-GAMMA, IL6 and TGF-BETA. These cytokines act on IDO1 via both inhibitory and activation protein interactions. This differentially expressed protein activating PI3K-AKT pathway is known to perform vital roles during leprosy [199].

### 3.2 Pathway analysis

*Mycobacterium Leprae*, the leprosy causing bacteria has shown high amount of activity in the five human signaling pathways i.e. mTOR signaling pathway, TLR signaling pathway, T-cell signaling pathway, NOTCH and MAPK signaling pathway [18,20,21,34,35]. So, we decided to do our pathway analysis with these five signaling pathways, the ultimate aim is to construct a global comprehensive leprosy based pathway. *M. leprae* like other bacteria has the ability to destruct the signaling interactions between the different pathway proteins and it has been seen to perform this mechanism in these five pathways in higher amount. From the databases, we got the basic structure of the pathway figures. However, databases consist of discrepancies; so, we searched for the interactions in literature articles and also included the new interactions which have been recently found. We tried to obtain more and more experimental evidences for the protein-protein interaction (PPI) from the leprosy based papers. We have mentioned “Yes” for the reference from a leprosy article and “No” if not from a leprosy article for each of the PPI.

The DEGs coding for 69 proteins have showed significant activity during leprosy, so, we hypothesized that they might be involved in one or more than one of the five pathways. We found out that a total of 48 proteins out of 69 were involved in one or more than one of the five pathways (Table 1).

**Table 1.**
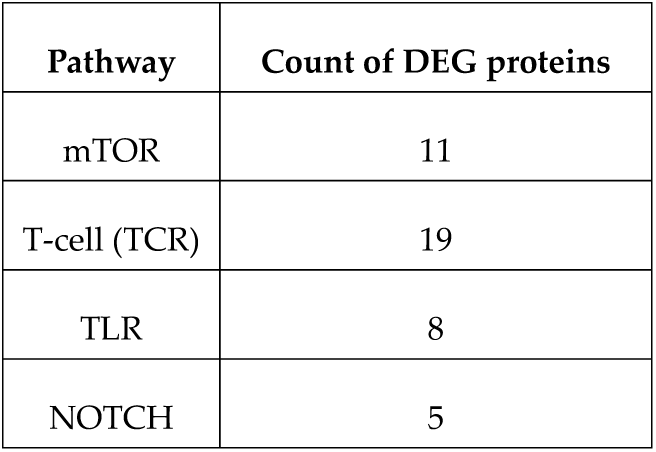
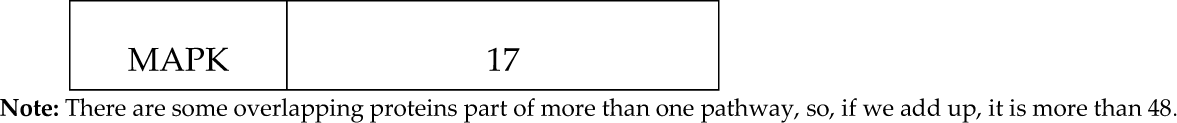
Number of DEGs involved in each of the five signaling pathways.

**Table 2.** Number of molecules and molecular interactions (PPIs) for each of the five pathways:

| Pathway component | mTOR | TLR | NOTCH | MAPK | T-cell |
| --- | --- | --- | --- | --- | --- |
| Total molecules | 44 | 68 | 59 | 88 | 78 |
| Total molecular interactions | 55 | 84 | 76 | 122 | 112 |

**Note:** Molecules comprise proteins, molecules such as Calcium, CpG islands and molecular interactions comprise majorly protein-protein interactions including host-pathogen interactions.

We first found out the interaction partners of our experimental host proteins to find that in which pathway they are involved using manual curation or literature survey. For some of the proteins, UniProt and BioGRID also gave us the interaction details and these databases can be trusted as they mention the literature article for their results [65,68]. For each of the interaction occurring for each of the five pathways including our experimental host proteins; we have made a one to one protein-protein interaction file with PubMed reference IDs for each of the interaction [64]. These files are given in Table. Once we have made the interaction file for the pathways, we constructed the five pathways, mTOR, TLR, TCR, NOTCH and MAPK signaling in the presence of M. leprae in pathway visualization software called CellDesigner [66]. CellDesigner uses the computing language SBML (Systems Biology Markup Langauage) to construct the pathway figures [67].

We are claiming that till date, our reconstructed pathway figures have maximum number of molecules/proteins and interactions occurring during leprosy through the five signaling pathways.

### 3.3. Pathway description of five leprosy-affected signaling pathways

#### 3.3.1 mTOR signaling pathway

The complement receptors CXCR1/2 and CXCR3 act as extracellular ligands for transmitting the signal in the macrophages through the mTOR pathway. **PPBP** physically interacting with CXCR1/2 and **CXCL9** and **CXCL10** with CXCL3 leads to signal transfer to GNAI [72–74]. PPBP is an up-regulated differentially expressed gene found from experimental studies of leprae infected armadillos. GNAI-SRC interaction leads to the activation of the PI3K-AKT1-mTOR cascade ultimately stimulating the transcription factors (TF’s) to produce the required output proteins [72,75–78]. RAS interacts with RAF for activation of ERK1/2 which activates TSC1/2 and this protein inhibits AKT1 [79,80]. *M. leprae* once it enters the cell is present inside a phagosome. There it releases the protein ESX-1 inside the host cell which causes the disintegration of the phagosome causing the release of the bacteria [24]. It has been recently identified that ManLam which is a regular surface component of Mycobacteria is found in *M. leprae* as LepLam [39]. This protein facilitates the important cross-talk interactions with the PI3K-AKT1-mTOR cascade by exploiting the Ca^2+^ signaling. PLC activation of PIP2 leads to the activation of IP3 and PI3K [81–83]. IP3 has been known to increase the Ca^2+^ concentration inside the cell which activates the CALMODULIN and after getting activated; it goes and binds PI3K [84–86]. LepLam inhibits the Ca^2+^ molecule, CALMODULIN and PI3K [25,39].

It has been clearly shown in a paper that PI3K interacts with EEA1 and **RAB7B** for the activation of autophagosome formation around the *M. leprae* bacteria [25,87]. RAB7B is also one of the differentially expressed proteins found from the experimental studies. Another protein LC3-II is hugely known for its role in autophagosome formation around the bacteria. **OPTN**, **MREG** and **P62** are the three proteins which by different pathways bind to LC3-II and recruit it to the *M. leprae* surface which then, there starts the autophagosome formation [26,88–92]. OPTN and MREG are again down-regulated differentially expressed gene. Some other relevant interactions have been included in our pathway consisting of mainly our differentially expressed genes. **GAS6** and **PLP1** (2 of our down-regulated gene) activates and physically interacts with AKT1 respectively [93,94]. Downstream effector of AKT1 is **FSTL1** is again a differentially expressed gene [95]. RAC1 protein binds with PARKIN which inhibits mTOR and activates AKT1 and RAC1 alone also interacts with mTOR [96–99]. RAC1 is inhibited by another down-regulated differentially expressed gene, **CNN1** [100]**. SSPN** or Sarcospan is another protein shown to activate AKT1 [101].

#### 3.3.2 TLR signaling pathway

The TLR receptors which are involved in the leprosy dynamics are TLR1/2, TLR4, TLR6 and TLR9 [32,102–104]. LIPOPROTEINS, the known ligand of TLR1/2 binds to it causing the interaction of TLR1/2 with MYD88:TIRAP complex [102,105]. This complex goes and physically interacts with IRAK, then IRAK binds with TRAF6, leading to the activation of TF, NF-kB [105–107]. This protein produces two major cytokines, IL10 and IL12 [34,108]. LPS, the ligand for TLR4 interacts with it making it function in the same way causing the activation of NF-kB [109]. The MYD88 independent pathway is also active by TLR4 where the receptor binds with TRAM:TRIF and goes on physically interacting with TAK1 [110,111]. TAK1 activates the transcription factor IRF3 to produce the output proteins, IFN-ALPHA and IFN-BETA [112–114]. Also, IRF3 has been seen to get activated by another pathway where complement proteins are involved. **C1R**, **SERPING1** and **C1S** are three complementary proteins which are part of our set of differentially expressed genes.

These three physically interact with complement system which goes and binds to complement receptor, CR [115]. CR is inhibited by *M. leprae* [25]. CR stimulates ERK1/2 and PI3K to activate IRF3 [116–119]. *M. leprae* bacteria produces many proteins which interact with different host proteins to create a favorable environment for their survival. Some of these proteins are LPRG, BFR and PGL1. PGL1 directly inhibits the receptors TLR1/2 and TLR4 [43]. BFR and LPRG are shown to physically interact and activate TLR1/2 respectively to cause the activation of the transcription factor, CEBP-BETA [28,29,120].

From various articles, we got the cascade which is involved in activation of CEBPBETA i.e. through SRC-PLC-GAMMA-CALCIUM-P38 [121–125]. CEBP-BETA is producing the protein ARGINASE-1 which cleaves and inhibits ARGININE. ARGININE activates INOS and this enzyme produces NO [120]. In this manner, bacteria decreases the production of NO by causing more and more production of ARGINASE-1 through CEBPBETA. VIT-D3 is also involved in the mTOR pathway. CEBP-BETA normally helps in the production of antimicrobial peptides through production of CYP27b1 which activates VIT-D3 [35,126]. VIT-D3 binds with VDR and binds with RXR to form active transcription factor in the nucleus: VDR:RXR [35]. This protein produces two antimicrobial peptides, CAMP and DEFB4A [126]. So, it is hypothesized that *M. leprae* actually uses the transcription factor (TF), CEBP-BETA for its own advantage to survive inside the host cell by working through the TLR1/2-CEBP-BETA cascade. Bacterial protein PGL1 also activates the host miRNA HSA-MIR-21, which inhibits the CYP27b1 [126]. Recently, a paper has clearly shown the bacteria interact with DC-SIGN receptor on the dendritic cells to activate RAF1 which activates P65 which again activates the transcription factor, NF-kB [31]. Another paper has depicted the *M. leprae* influence through the TLR6 pathway in Schwann cells (SC). TLR6 once sensed by *M. leprae* or when bacteria interacts with it, it stimulates the production of lipid droplets (LD) which consist of SC derived lipids and act as nutritional source for the bacteria. LD acts as catalytic site for PGE2 synthesis. PGE2 along with IL10 is responsible for the inhibition of IL12 and NO in SC’s [32]. **RAB7B**, one of our differentially expressed genes has been shown to perform lysosomal degradation of TLR4 [127].

*M. leprae* enters the cell in a phagosome where it releases another protein, DNA-HLP in the cytosol [104]. The DNA-HLP forms complex with host CpG which goes and interacts with endosomal TLR9. Another article has shown that TLR9 once it gets stimulated by DNA-HLP:CPG, it activates IGF1 which goes on activating the ARGINASE-1 [33]. Hence, this cascade can be considered as an alternative pathway taken by the bacteria through TLR9 to form ARGINASE-1 along with the TLR1/2 cascade discussed above [120]. **HBB** is an up-regulated differentially expressed gene which activates the NO [128]. The other proteins from our experiments, **TMIGD3**, **LIMCH1** and **CAVEOLIN-1** are also part of the TLR cascade. TMIGD3 inhibits the transcription factor, NF-kB, CAVEOLIN-1 carries out physical interaction with TRAF2 and TRAF2 then interacts with TRAM:TRIF [129–131]. LIMCH1 binds with TRIM25, which activates TRAF6 [132,133].

#### 3.3.3 NOTCH signaling pathway

NOTCH signaling pathway’s involvement has recently been found during leprosy by two papers [21,45]. Although, the investigation of this pathway during leprosy has been done less till now, but these two research articles clearly mention that this pathway is hugely responsible for the pathogenesis by *M. leprae*. Two NOTCH receptors (NOTCH1 and NOTCH2) are known to function during leprosy. JAGGED1 is the extracellular ligand of the NOTCH pathway which interacts with these two NOTCH receptors to start the signaling [134,135]. Once they have interacted, TACE enzyme cleaves the two NOTCH receptors, NOTCH1 into NECD1 and NEXT1 and NOTCH2 into NECD2 and NEXT2 [136]. The next complex, GAMMA-SECRETASE cleaves the NEXT1 and NEXT2 into NICD1 and NICD2, which after translocating to the nucleus, shown as NUC-NICD1 and NUC-NICD2, act as transcriptional co-activator and co-repressor [44,137]. GAMMA-SECRETASE is a complex of many proteins, PRESENELIN1, NICASTRIN, APH1 and PEN2 [138]. TGF-BETA also stimulates JAGGED1 [139]. TLR1/2 has recently been identified as a receptor in NOTCH pathway [21]. *M. leprae* proteins BFR, LPRG, PIM2 and PGL1 signal through this receptor to disintegrate the host interactions using NOTCH signaling [28–30,41,140,141]. BFR physically interacts with TLR1/2, LPRG and PIM2 activate the TLR1/2, PGL1 inhibits TLR1/2 [28,29,41,43]. TLR1/2 physically interacts with PI3K which activates AKT [41,142]. AKT then activates ERK1/2 and ERK1/2 stimulates NF-kB [41]. NFkB produces the cytokines MMP9, IL12, IL10 and COX2 [34,41,108]. TLR1/2 is also signaled through MYD88:TIRAP and this complex binds IRAK which then binds TRAF6 for the activation of NF-kB [105,106,143,144]. NUC-NICD1 in the nucleus physically interacts with COA complex (MAML, EP300,SKIP and HAT) and also with COR complex (HDAC, SMRT, CIR, SIN3A, SAP30) [44,145–150]. NUC-NICD1 also binds SMAD3 [151]. All these functionalities are performed by NUC-NICD1 for regulating the activity of the most important transcription factor of this pathway: CSL. COA activates and COR represses CSL [44]. Similarly, NUC-NICD2 interacts with COA and COR for regulation of CSL through activation and repression of it respectively [44]. NUC-NICD1 and NUC-NICD2 also directly activate CSL [44]. CSL is responsible for the output proteins, HES1, SOCS3, COX2, ACTA2 and MMP9 [41,45,139,152]. NUC-NICD2 activates another transcription factor, YY1 which produces MYC [153]. ACTA2 is one of our differentially expressed protein. We have also got a negative feedback loop in the form of HES1 inhibiting PTEN which inhibits the intermediate protein, PI3K [142]. SLC25A4, a differentially expressed protein binds with transcription factor, TCF3 which produces HES1 [154,155]. SSPN (Sarcospan) activates AKT which produces the protein, FSTL1 [95,101]. Both of these are differentially expressed protein. The fifth differentially expressed gene part of this pathway is PLSCR4 which physically interacts with TLR1/2 [156].

#### 3.3.4 MAPK signaling pathway

EGFR is the only signaling receptor in leprosy affected MAPK signaling pathway [157–159]. It activates the GRB2:SOS complex leading to the activation of PI3K through RAS and via PI3K-AKT pathway, ERK1/2 is stimulated. ERK1/2 is responsible for mediating any signaling interactions in this pathway including activation of the transcription factor; NF-kB [41,160–162]. This protein produces GM-CSF, IL2, VEGF, BCL-XL and IL6 in this network [161,163–166]. RAS also activates the complex RAC:CDC42 which activates some intermediate proteins to finally activate P38 which activates another transcription factor of MAPK pathway; AP1 [167–169]. AP1 produces IL2, IL4, IL5, P19, CYCLIND1, CYCLIN-A and GM-CSF [170–175]. EGF protein acts as the main signal which activates the receptor EGFR in this pathway [157,159]. Many differentially expressed proteins are part of this pathway and one of them **LAMA2** is actually playing a huge role in this cascade. *M. leprae* releases the protein, PGL-1 and ML-LBP21 in the extracellular space and they interact with **LAMA2** [30,38,176]. **LAMA2** then binds to EGFR enhancing the MAPK signaling transmission [7]. *M. leprae* when it comes inside the cell, its coat protein LEPLAM also activates ERK1/2 [39,177]. Another bacterial protein released inside the cell, MURAMYL-DIPEPTIDE physically interacts with NOD2 which then binds RIP2 and it then again activates NF-kB [40,178]. *M. leprae* also releases PIM2 which activates PI3K in the pathway [41]. ETS and CREB are other transcription factors which are getting activated by ERK1/2 and are also producing several output proteins in this network including FASL, BCL2, PDGF, IL3, GM-CSF and CYCLIN-D1 [164,175,179]. Differentially expressed proteins **CAVEOLIN-1** and **HBB** are very much involved in this pathway. **CAVEOLIN-1** physically interacts with EGFR and **HBB** which is the up-regulated differentially expressed protein (dark blue) is binding with another MAPK protein, MAPK6 [180,181].

#### 3.3.5 T-cell signaling pathway

The antigen of *Mycobacterium leprae* is presented by MHC-CLASS-II molecule to the antigen presenting cells (APC’s) and other immune cells [29,36]. This leads to the activation of the T-cell. The main receptors involved in this pathway are TCR:CD3 complex and CD4 along with CD28 and CTLA4 which receives the co-stimulatory signal from the APC’s [29,36,182,183]. The TCR complex activates ZAP70 which activates LAT [184,185]. LAT then activates PLC-GAMMA which interacts with PIP2 to give DAG and IP3 [186–189]. DAG activates PKC-THETA which finally activates one of the most important transcription factors of this network; NF-kB [190,191]. NF-kB produces important output proteins of this pathway which are IL2, IL6, GM-CSF, IFN-GAMMA and IL10 [37,108,161,192,193]. NFAT is another transcription factor which is activated by GSK3-BETA through the PI3K-AKT axis which produces IL2, IL3, IL4, IL5 and TNF-ALPHA [171,194–198].

Another receptor TNFAR through an alternate route activates TRADD which activates TRAF2 [200]. TRAF2 stimulates ASK1 which activates MKK [201,202]. MKK then activates JUN and P38 [202,203]. P38 activates another transcription factor, AP1 [169]. AP1 is responsible for producing the proteins, IL2, IL3, IL4, IL5, IL6, TGF-BETA and GM-CSF [171,172,175,204–206]. Many differentially expressed proteins are part of T-cell network. The major ones which are regulating the cascade to a huge extent are CAVEOLIN-1, IDO1 and SSPN. SSPN physically interacts with AKT which is regulating this network at many crucial points [101]. CAVEOLIN-1 performs physical interaction with TRAF2 and activates ASK1 and signal flow through this protein leads to the activation of P38 protein [130]. IDO1 has been seen to activate directly NF-kB which makes it a significant protein of this pathway [207]. Many inhibitory interactions are happening too which is helping in the functioning of this pathway. Calcium channel CRAC in the membrane causes the calcium uptake which activates CALCINEURIN [208,209]. This protein inhibits NFAT [210]. NF-kB is also inhibited by differential expressed TMIGD3 along with IKB-ALPHA and IKB-BETA [129,211,212]. These two proteins are activated by IKK-BETA which is getting stimulated from the TNFAR receptor signaling axis [211–213]. The two *M. leprae* proteins are LPRG and BFR which are released by the bacteria and they are presented by MHCCLASS-II to the APC’s [29,214]. APC then activates the T-cell signaling network. TNF-ALPHA and IL2 are the output proteins and after they are produced they again come out of the cell and bind to TNFAR and IL2R respectively in the T-cell membrane creating a positive feedback loop in this signaling cascade [215,216].

### 3.4. Gene ontology (GO) characteristics of differentially expressed host proteins

We looked for Gene Ontology (GO) Biological Process terms for each of these 69 genes using the Uniprot Interface [68,69]. We chose to focus on two generalized biological functions associated with the disease of leprosy: Immunological and Neurological. Immunological functions depict that the genes are involved in some kind of immunity against the bacteria and neurological functions as many leprosy cases have been associated with neuropathies. We created 3 sets of proteins: 1) having immunological terms 2) having neurological terms and 3) having both immunological and neurological terms. These terms were not kept very specific but general e.g. terms like MHC-II complex binding and astrocyte development also have been considered under immunological and neurological terms respectively. In this way, we obtained 49 proteins out of 69 which in total formed the above mentioned 3 sets.

**Figure 7.**
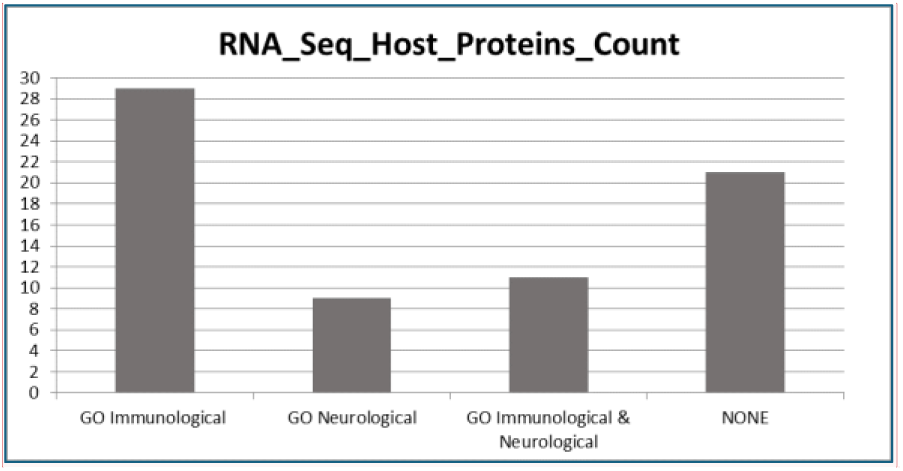
GO classification of differentially expressed host proteins. Out of 49 proteins, 29 proteins consisted of GO Immunological terms, 9 proteins consisted of GO Neurological terms and 11 proteins consisted of both GO Immunological and Neurological terms [69]. Hence, 49 proteins out of 69 depicts that 71% of the proteins which were experimentally found are predicted to be involved with the disease of leprosy in some or the other way. However, there were also 20 proteins which did not show either GO Immunological or Neurological terms (NONE).

### 3.5. Identifying biomarkers in M. leprae-affected signaling pathways using graph theory

The values for each of the centrality parameters mentioned in Section 2.6 were computed for each of the proteins of the signaling cascade [47,49,54]. Then, we extracted the proteins which were showing values above average in each of the parameters. Among those, we observed that some proteins had values which were reasonably high compared to others. In the upcoming bar plots (Figure 8-12), we show those proteins and explain their potential functions when *M. leprae* infects a host cell in Table 3-7. Interestingly, in each of the centrality parameters, a differentially expressed protein from the wet lab experiments (Section 3.1) came up indicating our graph theory results are reliable. Degree connections are totaled in the Total Degree (In-degree + Out-degree).

#### a) mTOR signaling pathway

**Figure 8.**
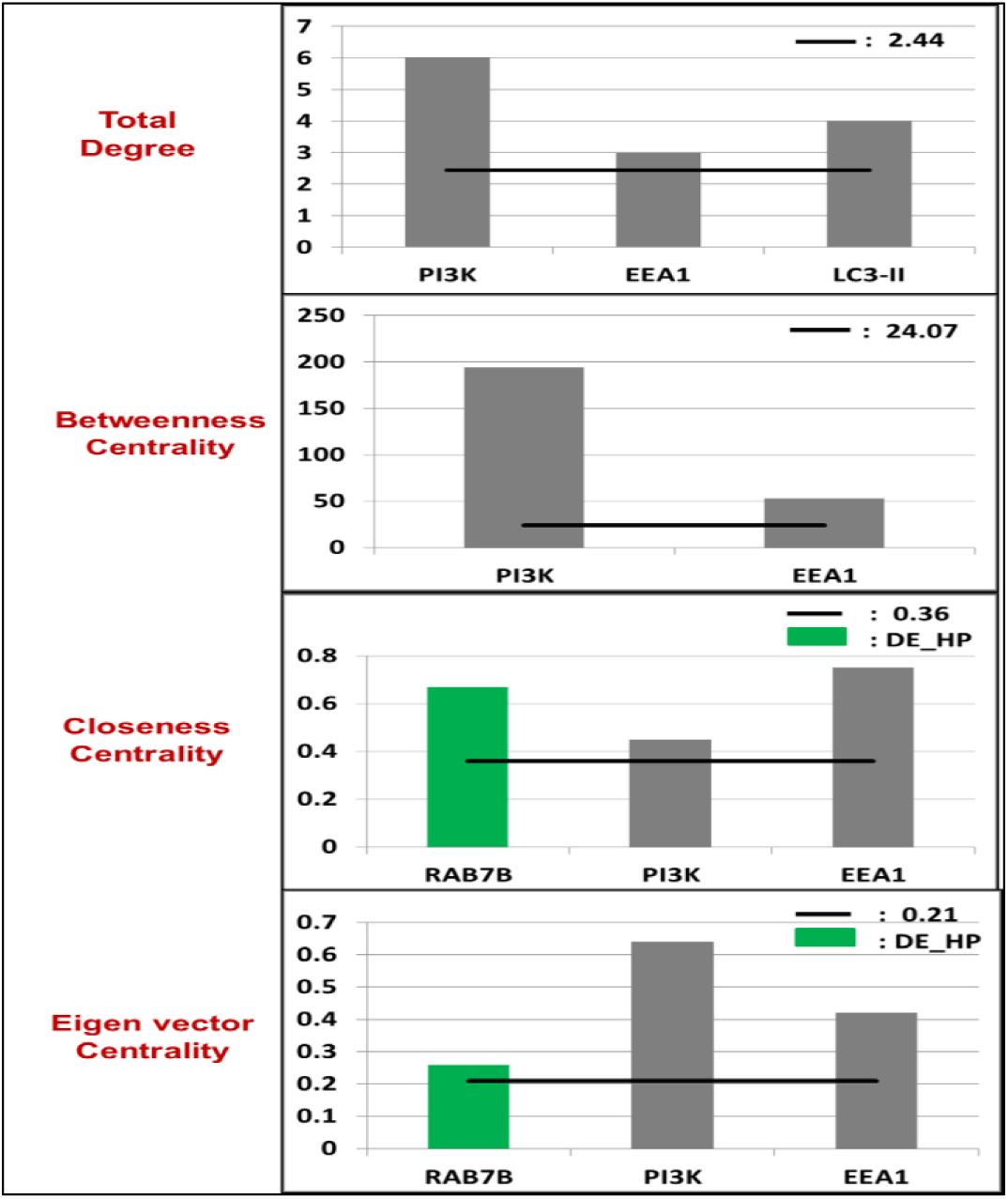
Potential biomarkers in mTOR signaling pathway proteins. Differentially Expressed Host Protein (DE_HP) is shown in green.

**Table 3.** How mTOR pathway proteins can affect leprosy dynamics in humans.

| Protein | Involvement in leprosy | Reference |
| --- | --- | --- |
| LC3-II | Interact with many proteins including 2 DE_HPs: Optineurin and Melanoregulin to kill *M. leprae* by forming autophagosomes. | [26,27,88,217] |
| PI3K | Binds AKT and mTOR to function during chronic neuropathic pain: a disorder commonly observed in leprosy patients. | [25,76,218,219] |
| EEA1 | Forms complex with RAB7B and PI3K to form autophagosomes around Mycobacteria to kill it. | [25,87] |
| RAB7B | A DE_HP binding EEA1 to activate autophagosome formation around *M. leprae* and kill it. | [25,87] |

#### b) TLR signaling pathway

**Figure 9.**
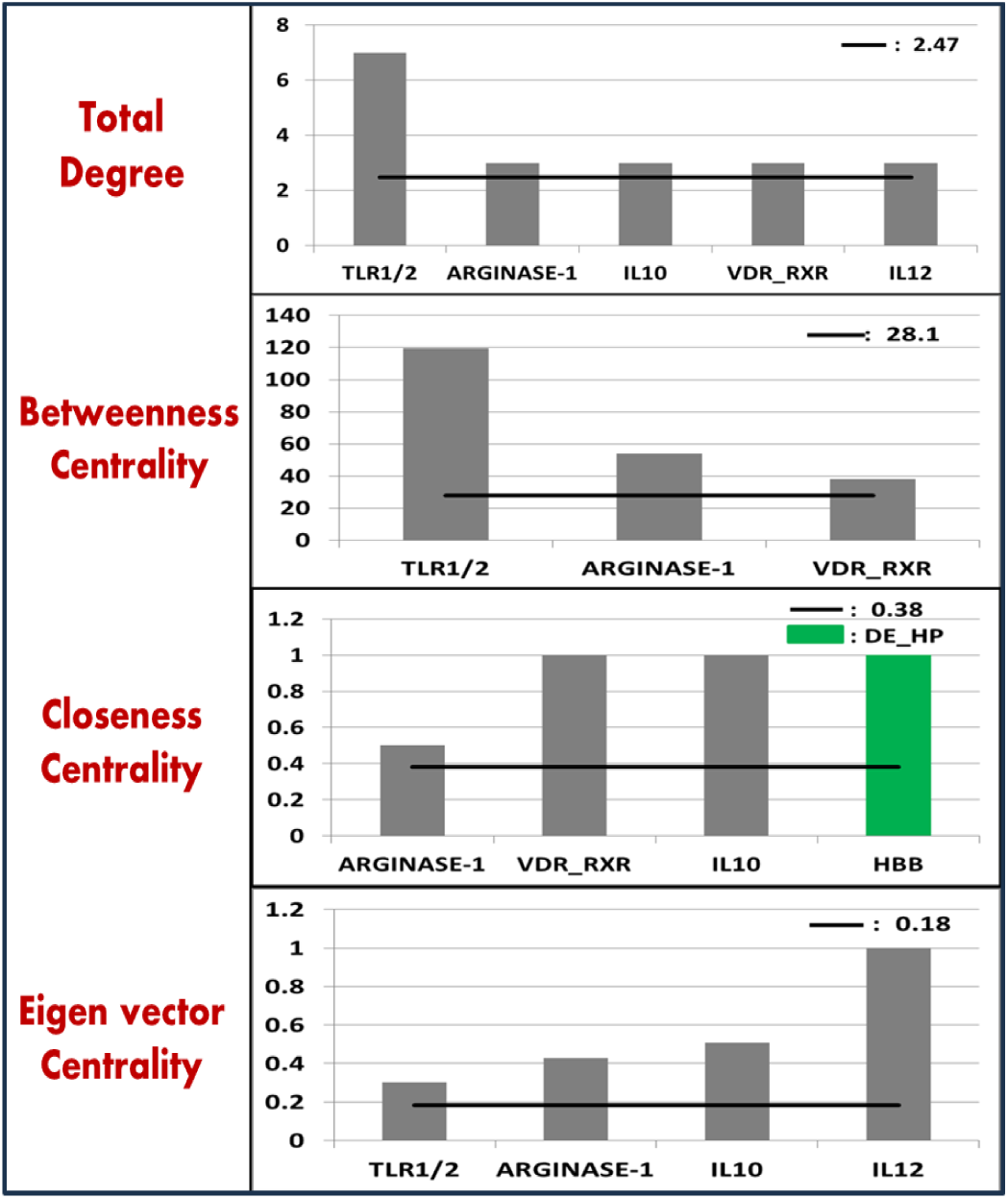
Host proteins of TLR signaling pathway predicted as biomarkers to detect leprosy early in patients. Proteins with differential expressions are shown in green (DE_HP).

**Table 4.** Function of TLR pathway proteins in M. leprae infected host cells.

| Protein | Involvement in leprosy | Reference |
| --- | --- | --- |
| TLR 1/TLR 2 | Interact with multiple *M. leprae* proteins. The bacteria ensure its survival via binding these receptors to activate TF CEBP_BETA forming ARGINASE-1. | [28,29,43,105,108,220] |
| ARGINASE-1 | Once *M. leprae* activates this enzyme via TLR signaling, it cleaves arginine lessening production of Nitric oxide (NO) as this molecule prevents bacterial invasion of the host cell. | [120] |
| IL10 | Anti-inflammatory cytokine whose expression usually increases in lepromatous leprosy patients. | [221,222] |
| IL12 | Inflammatory cytokine helping in activating defense mechanisms by the host cell against *M. leprae*. | [223,224] |
| VDR-RXR | A TF producing antimicrobial peptides CAMP and DEFB4A to reduce *M. leprae* effects on the host. | [35,126] |
| HBB | This DE_HP produces NO and thus, mediates *M. leprae* killing. | [128] |

#### c) NOTCH signaling pathway

**Figure 10.**
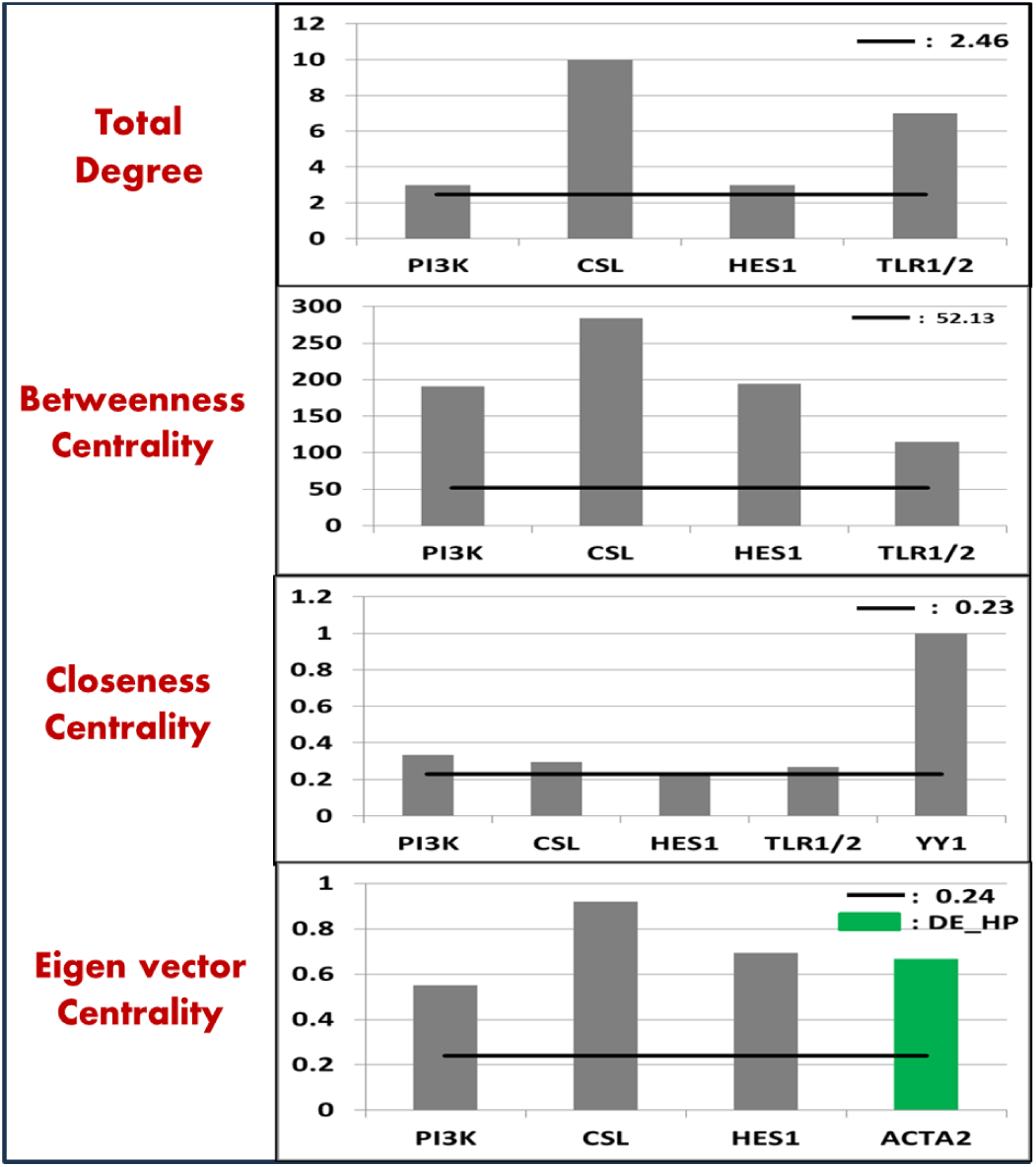
Proteins identified as leprosy regulators during NOTCH signaling. ACTA2 in green is the Differentially Expressed Host Protein (DE_HP).

**Table 5.**
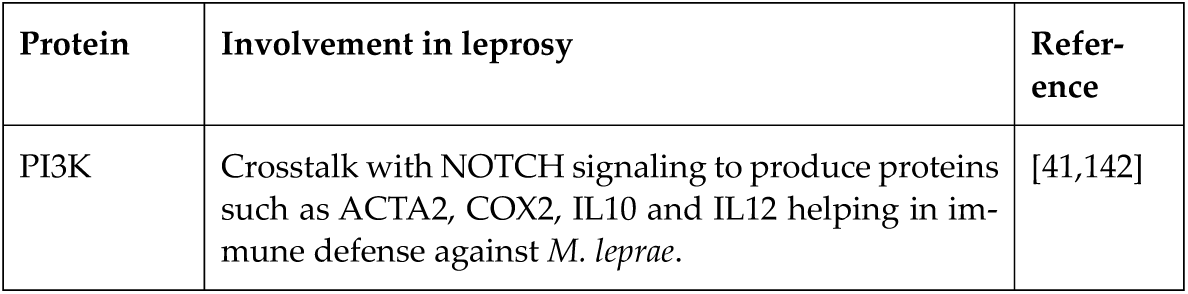
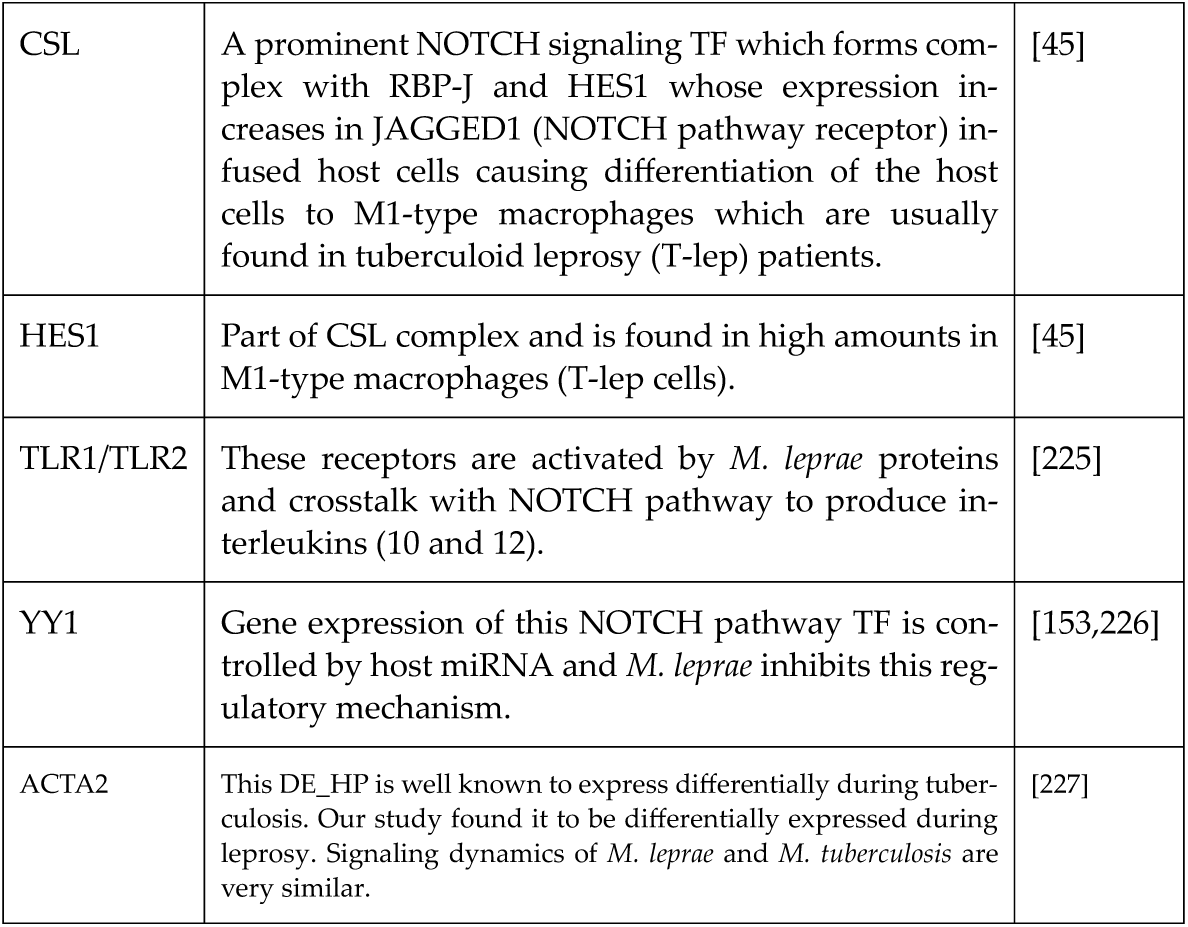
Proteins of NOTCH pathway which help in *M. leprae* pathogenesis.

| Protein | Involvement in leprosy | Reference |
| --- | --- | --- |
| PI3K | Crosstalk with NOTCH signaling to produce proteins such as ACTA2, COX2, IL10 and IL12 helping in immune defense against *M. leprae*. | [41,142] |
| CSL | A prominent NOTCH signaling TF which forms complex with RBP-J and HES1 whose expression increases in JAGGED1 (NOTCH pathway receptor) infused host cells causing differentiation of the host cells to M1-type macrophages which are usually found in tuberculoid leprosy (T-lep) patients. | [45] |
| HES1 | Part of CSL complex and is found in high amounts in M1-type macrophages (T-lep cells). | [45] |
| TLR1/TLR2 | These receptors are activated by *M. leprae* proteins and crosstalk with NOTCH pathway to produce interleukins (10 and 12). | [225] |
| YY1 | Gene expression of this NOTCH pathway TF is controlled by host miRNA and *M. leprae* inhibits this regulatory mechanism. | [153,226] |
| ACTA2 | This DE_HP is well known to express differentially during tuberculosis. Our study found it to be differentially expressed during leprosy. Signaling dynamics of *M. leprae* and *M. tuberculosis* are very similar. | [227] |

#### d) MAPK signaling pathway

**Figure 11.**
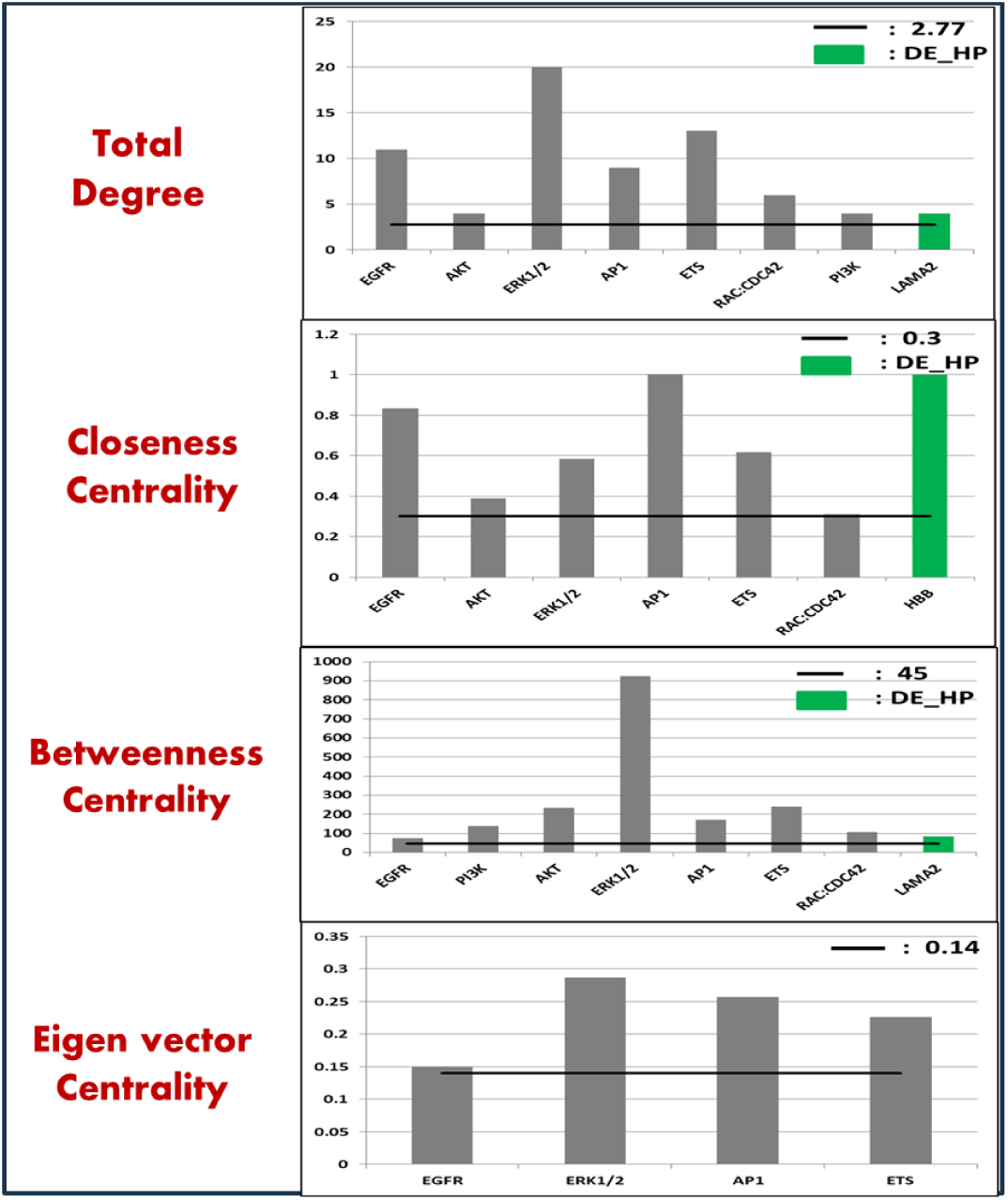
Multiple proteins of MAPK signaling pathway actively functions when *M. leprae* hijacks host cell. 2 DE_HPs; LAMA2 and HBB are part of this signaling pathway (green).

**Table 6.**
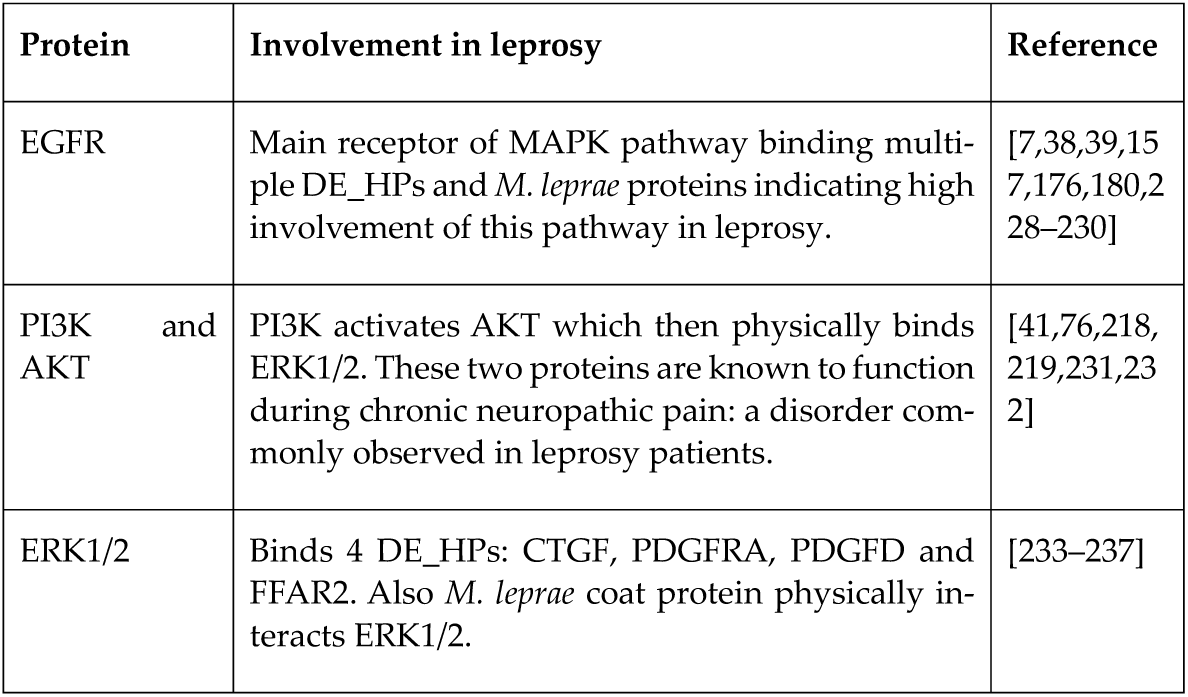
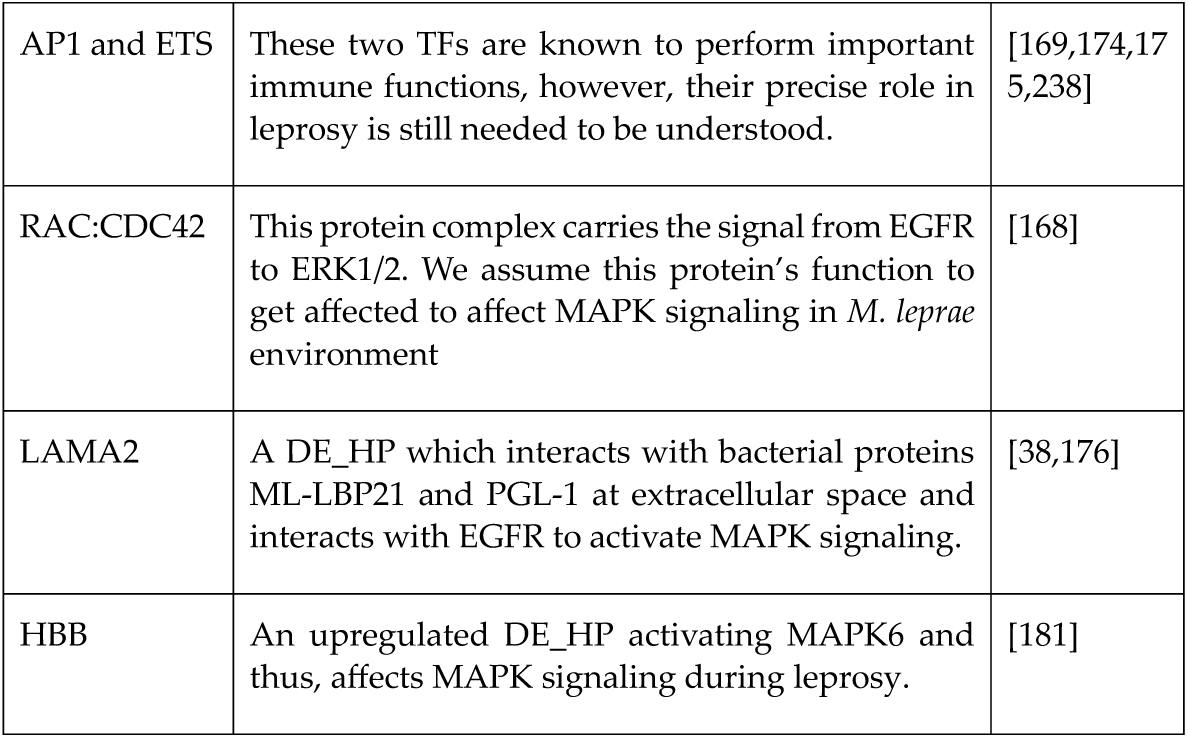
MAPK signaling pathway function during leprosy via a high number of proteins.

#### e) T-cell signaling pathway

**Figure 12.**
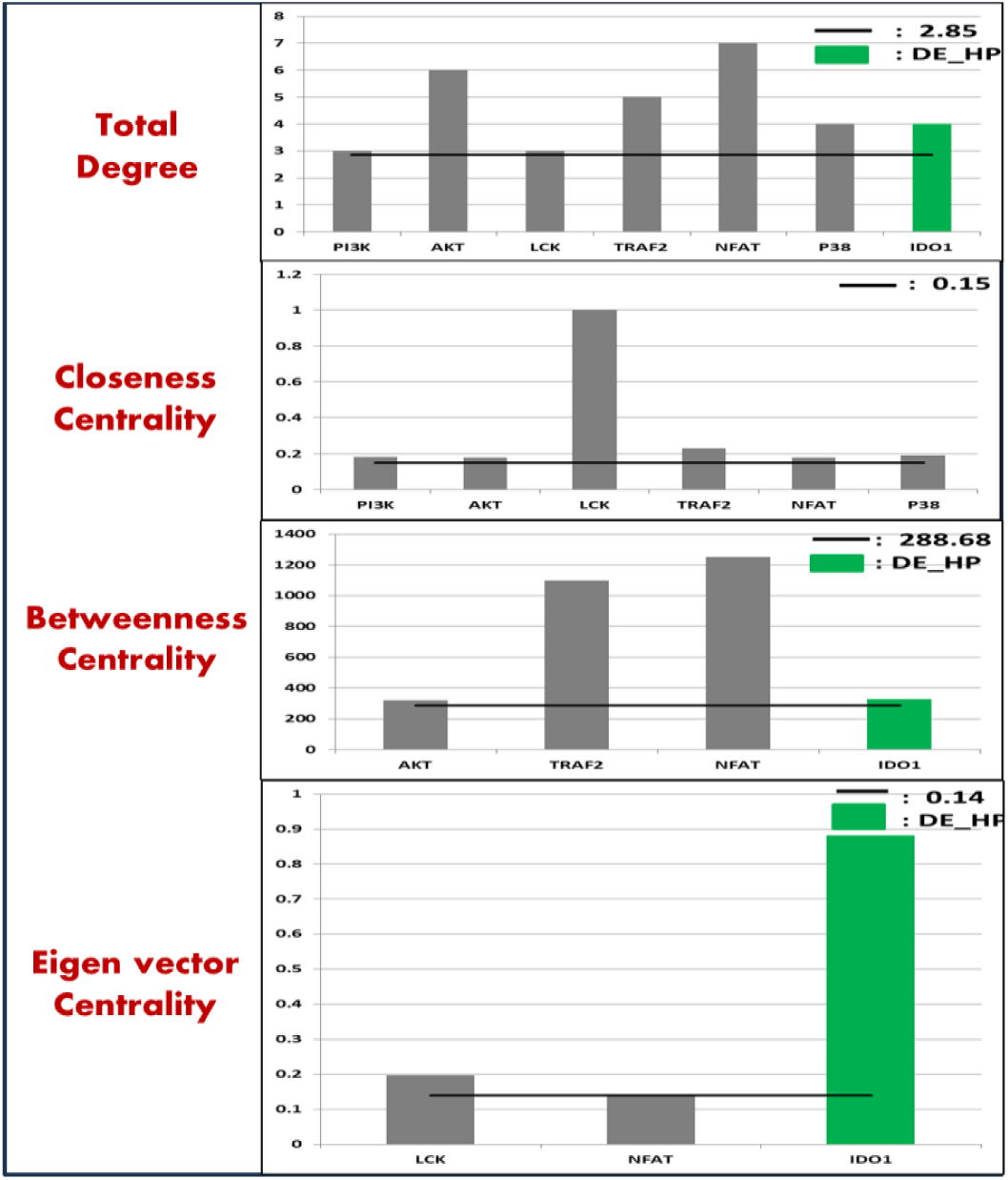
Different types of proteins in T-cell are active during leprosy including transcription factors TRAF2 and NFAT and DE_HP IDO1 (green) [199].

**Table 7.** T-cell proteins with immune defense functions against a foreign bacteria.

| Protein | Involvement in leprosy | Reference |
| --- | --- | --- |
| PI3K | Gets activated by CD28 and produce crucial output proteins via TF NFAT and NF-κB during leprosy. | [194,232,239,240] |
| AKT | Downstream regulator of PI3K leading to activation of NFAT and NF-κB. | [195,198,241] |
| LCK | T-cell receptors ZAP70 and CD4 activate this protein. | [242–244] |
| TRAF2 | This protein facilitates the activation of another TF AP1 in T-cell pathway during *M. leprae* hijacking of host environment. | [201,202,245] |
| NFAT | TF producing crucial output proteins such as interleukins, TNF-ALPHA. | [125,171,246,247] |
| P38 | Another protein activating crucial T-cell pathway proteins when a foreign pathogen attacks it such as AP1, FOS, CREB etc. | [248–251] |
| IDO1 | A DE_HP activated by NF-κB. Also itself activates TGF_β and IFN_γ during T-cell pathway. Already studied in context to leprosy. | [207] |

## 4. Discussion

Leprosy prevalence has declined significantly all over the world in last 3 decades. However, in the past decade, the new case detection has remained stagnantly high at over 200,000 cases annually worldwide. To understand the transmission dynamics of this disease, it is important to develop tools for early detection of the disease. For this, the host or pathogen based biomarkers are to be identified. For early detection when pathogen number is below detection limits of most of the available assays, it is important to identify host-biomarkers by understanding the biological mechanisms upon *M. leprae* infection which lead to disease progression in the susceptible hosts while the resistant ones are able to control the infection. Therefore it is important to understand the associated signalling pathways in more detailed manner. Not only *M. leprae*, *Leishmania major* and *M. tuberculosis* also are known to manipulate the host environment to their advantage by affecting these signaling pathways for their advantage [56,252]. It is obvious that bacteria is not targeting only one protein or one complex or one pathway only at a time. Therefore, we need to investigate the mechanisms of these bacteria through a systems biology approach which will be helpful to comprehend the intricate processes of *M. leprae* [15]. Once we understand the signaling routes taken by the bacteria in our body, we can select the biomarkers and check their RNA expression by qPCR or RNA sequencing analysis [253,254]. The goal of this work is not to find therapeutic drug targets for this disease as both multi- and pauci-bacillary leprosy can be cured nowadays with antibiotic therapy but having information of the “leprosy-affecting” host proteins will help in diagnosing the disease early in humans before the pauci-bacillary progress to multi-bacillary leprosy in them. Till now, no one has found, even the crucial regulatory proteins which form essential part in leprosy dynamics. Biomarker identification or checking the RNA expression will validate that these host proteins are exclusively expressed during leprosy [255]. Definitely, after that, it can be directed towards wide range of therapeutic or biomedical applications. In this study, five major signaling pathways (mTOR, TLR, NOTCH, T-cell and MAPK) have been investigated in reference to leprosy progression using the available literature [18,21,35,161,240]. For this, the signaling pathways dysregulated upon leprosy progression were drawn in the pathway visualization software CellDesigner to depict the biological interactions clearly [66]. Not only we incorporated the bacterial processes which *M. leprae* performs to invade the host cells, but, our model has the human homologs of up-regulated and down-regulated host proteins which were differentially expressed in the RNA sequencing of leprosy affected armadillos. Thus, this study is a first to provide a comprehensive reference to the human signaling pathways affected in leprosy using systems biology approach and provides suitable literature reference [64]. These interaction files are given in the supplementary Table S2. We have performed validation of the 69 differentially expressed proteins in two ways. First, the literature survey gave us 48 proteins which are forming part of our leprosy pathway models. Second, the database survey gave us 49 proteins (Figure 7) which are involved in leprosy in some or other way as they showed GO biological process immunological and neurological terms in UNIPROT [68,69]. The overlap of these two sets gave us 42 proteins (Figure 13, Table S3).

**Figure 13.**
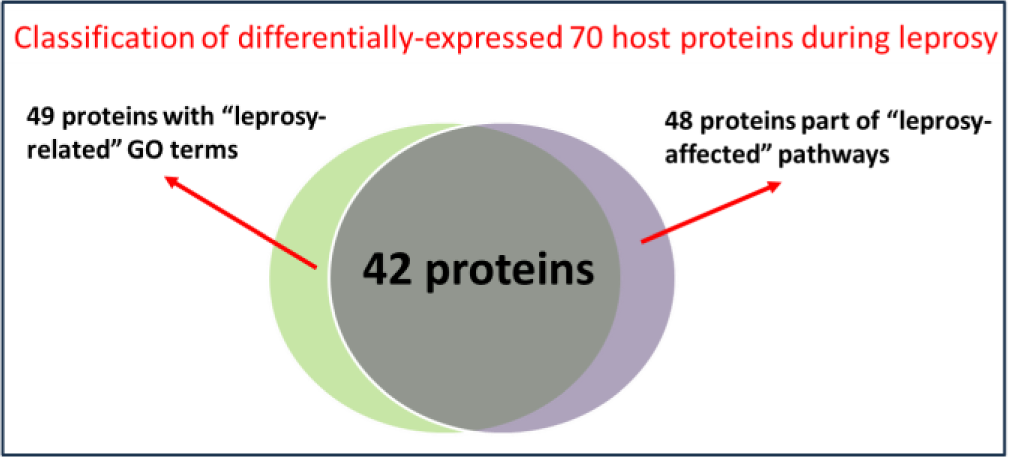
Our study verified experimentally identified proteins (42 proteins out of 69) are highly likely to play crucial roles when *M. leprae* hijacks the human cell.

These proteins were identified from RNA sequencing results hence, we have experimental basis of their involvement/association in leprosy. But directly choosing them as host biomarkers might not help as mentioned above, *M. leprae* is targeting many proteins at a single time and we have to understand the upstream and downstream regulators of these differential expressed proteins and also of the bacteria [25]. We can only do that when we draw and visualize the signaling pathways and then look at the paths followed by the bacteria. And as we expected, when we did the analysis, we got wide range of host proteins which can be involved in leprosy. However, since proteins were identified based on showing altered RNA expressions in uninfected vs *M. leprae* infected armadillos, there is a possibility that the differentially expressed RNA may code same proteins of different functions/structures [256]. Tackling this limitation of proteoforms will be a bit out of scope for this study. But this study very well captures another crucial aspect of proteoform complexity: proteins rarely function alone but usually interact with other proteins [257]. We applied graph theory analysis on five different protein-protein interaction networks: signaling pathways. The centrality measures in graph theory which considers count of PPIs to find the hub proteins in a network were predicted as important proteins influencing the leprosy dynamics [49,53]. Accordingly, we have chosen biomarkers from each of the pathways from the ones which are showing above average values in the graph theory centrality parameters. The significance for each of the biomarkers has been mentioned above.

Our graph theory analysis has given best results for the NOTCH signaling pathway. There are very few papers which have shown till now the involvement of leprosy through this pathway. But those 2 papers acted as solid reference to convince us to incorporate this pathway into our analysis [21,45]. One of the papers have shown that endothelial cells when infused with the NOTCH ligand Jag1 and IFN-GAMMA, they differentiate the monocytes to become M1 type macrophages which are usually found in tuberculoid leprosy (pauci-bacillary) patients and when it is not infused with JAG1, the monocytes develop to lepromatous (or multi-bacillary) leprosy [45]. Also, the endothelial cells showed up-regulation of RBP-J, SOCS3 and HES1. RBP-J also known as CSL is the main transcription factor and it has showed good centrality values making it an effective biomarker [145]. In the same way, HES1 has also been chosen as a host biomarker due to its higher centrality values. TLR1/2 is another biomarker of NOTCH pathway and it induces the lipopeptides during leprosy [21]. YY1 is another biomarker which is attacked by host miRNA during leprosy [226]. Therefore, all the biomarkers have good references to call them as effective biomarkers. Along with that, the added advantage is NOTCH is a small signaling pathway and due to the low number of reports of its involvement during leprosy, we feel it will be a good area to start experimental studies with this pathway. PI3K has proved to be a global biomarker as it has been chosen as biomarker for each of our pathways except TLR pathway. It has showed higher centrality values in all the 4 pathways, mTOR, NOTCH, T-cell and MAPK. In mTOR signaling pathway, PI3K is inhibited by LEPLAM, the coat protein of *M. leprae* [25]. It activates AKT to produce output proteins and binds with EEA1 and RAB7B to form the autophagosme around the bacteria to kill it [25,76]. In NOTCH pathway, it is activating AKT which is stimulating NF-kB through ERK1/2 to produce crucial NOTCH output proteins [41,142]. In T-cell pathway, it is mediating the activation of both important transcription factors, NFAT and NF-kB [76,190,194,232]. PI3K is responsible for the ERK1/2 activation through AKT and PIP2 in MAPK signaling pathway [76,232].

The biomarkers which we have tried to find are computationally predicted. Therefore, experimental studies for the NOTCH biomarkers need to be done in future for validation. Apart from NOTCH pathway proteins, we propose 3 host proteins: LC3-II, PI3K and IDO1 whose abnormal expression in a patient is highly likely to be a signature of *M. leprae* infection. LC3-II kills *M. leprae* by forming complexes with multiple DEGs [27,88]. PI3K shows high values when graph theory applied on 4 networks indicating its involvement in high and crucial protein-protein interactions during leprosy [8]. IDO1 has already been also reported in previous studies in leprosy patient samples as a potential biomarker [199], however, our study provides another evidence for its role in disease progression and disease pathogenesis. Further experimental validation of these biomarkers using qPCR and RNA sequencing in human samples should be the next step towards clinical trials for this mycobacterial disease [253,254].

## Data availability

RNAseq transcriptomic data available at NCBI with Bioproject ID PRJNA1204495 (Table S1).

## Acknowledgements

Authors are thankful to the NHDP Baton Rouge staff particularly Vilma Marks, Heidi Zhang, and Roena Stevenson for the armadillo experiments. Authors are also grateful for the funding support for this work from the Department of Biotechnology, Indian Council of Medical Research (Government of India), Turing Foundation and the Leprosy Research Initiative, Netherlands (Project number 704.15.59; 708.20.09),

R2STOP Canada and an Interagency Agreement [AAI15006] between the Health Resources and Services Administration and the National Institute of Allergy and Infectious Diseases.

## Notes

### Competing Interest Statement

The authors have declared no competing interest.

